# Strain-Specific Tropism and Transcriptional Responses of Enterovirus D68 Infection in Human Spinal Cord Organoids

**DOI:** 10.1101/2025.06.27.661907

**Authors:** Nathânia Dábilla, Sarah Maya, Colton McNinch, Taylor Eddens, Patrick T. Dolan, Megan Culler Freeman

## Abstract

The mechanisms by which *Enterovirus* D-68 (EV-D68) infection leads to acute flaccid myelitis (AFM), a severe neurological condition characterized by sudden muscle weakness and paralysis, remain poorly understood. To investigate the cellular tropism and infection dynamics of EV-D68, we profiled naive and EV-D68-infected human spinal cord organoids (hSCOs) derived from induced pluripotent stem cells (iPSCs) using single-cell RNA sequencing (scRNA-seq). Examining the cellular composition of healthy hSCOs, we found that hSCOs comprise diverse cell types, including neurons, astrocytes, oligodendrocyte progenitor cells (OPCs), and multipotent glial progenitor cells (mGPCs). Upon infection with two EV-D68 strains, *US/IL/14-18952* (a B2 strain) and *US/MA/18-23089* (a B3 strain), we observed distinct viral tropism and host transcriptional responses. Notably, *US/IL/14-18952* showed a significant preference for neurons, while *US/MA/18-23089* exhibited higher rates of infection in cycling astrocytes and OPCs. These findings provide novel insights into the host cell tropism of EV-D68 in the spinal cord, offering insight into the potential mechanisms underlying AFM pathogenesis. Understanding the dynamics of infection at single-cell resolution will inform future therapeutic strategies aimed at mitigating the neurological impact of enteroviral infections.

## Introduction

Acute flaccid myelitis (AFM) is a polio-like illness characterized by muscle weakness and paralysis, primarily affecting children [1,2]. Increased cases of AFM suspected to be due to enteroviral infection were first recorded in 2014 [3–5] and many of such cases have been associated with *Enterovirus* D68 (Taxonomy: *Enterovirus deconjuncti*), or EV-D68 [6–9]. Subsequently, EV-D68 and AFM had coinciding biennial outbreaks from 2014 to 2018 [10], with the 2018 AFM outbreak associated with nearly twice as many confirmed AFM cases compared to 2014 [2]. Another spike of cases was expected in 2020, but transmission was likely impeded by isolation policies during the SARS-CoV-2 pandemic [11]. EV-D68 had an additional outbreak in 2022, but AFM cases did not increase as expected [2,12].

The mechanism by which enteroviral infection contributes to the development of AFM is unknown. Previous studies have suggested both direct damage to spinal cord neurons after viral infection, and subsequent cytotoxic T-cell responses to infected neurons both contribute to disease [9,13]. One fundamental question relevant to EV-D68 pathogenesis is the cell types that contribute to virus replication and production in the CNS. Studies in multiple model systems have demonstrated that EV-D68 can target and replicate in neurons [14–18]. Astrocyte infection has also been identified during EV-D68 infection of murine brain slice cultures and primary human astrocytes *in vitro* [19,20].

We have previously shown that contemporary strains of EV-D68 can replicate in an induced pluripotent stem cell (iPSC)-derived human spinal cord organoid (hSCO) model, which provides a valuable human-derived, multicellular model in which to explore EV-D68 pathogenesis [21]. Analysis of marker gene expression suggests hSCOs comprise multiple cell types, including neurons and glial cells, but the specific cell types present, and which contribute to enteroviral infection in the hSCO model are unknown [21].

To better define the cell types infected by EV-D68, and to identify potential strain-specific differences in cell tropism and pathogenesis, we infected hSCOs with two contemporary strains of EV-D68 associated with AFM, US/IL/14-18952 (18952) and US/MA/18-23089 (23089). These strains are genetically distinct (18952 is a B2 strain and 23089 is a B3 strain) and represent the circulating strains from the 2014 and 2018 outbreak years [22–24] (Fig 1). Using single-cell RNA sequencing (scRNAseq), we captured host and viral transcripts within different cell types and subtypes. Our analysis revealed the complex cellular composition of hSCOs, which includes neuronal and glial cell lineages. Analyzing viral transcript abundance within these cell populations demonstrated that the EV-D68 strains exhibit markedly different tropisms. Although EV-D68 18952 was primarily associated with neuronal infection, 23089 exhibited a preference for cycling astrocytes. Subsequent analysis of host cell transcriptional responses in both infected and bystander cell populations revealed further differences between these strains. Together, these findings clarify the shifting cell tropism of EV-D68 strains and provide insight into the mechanism of AFM pathogenesis in a highly relevant human model system.

**Figure 1.**
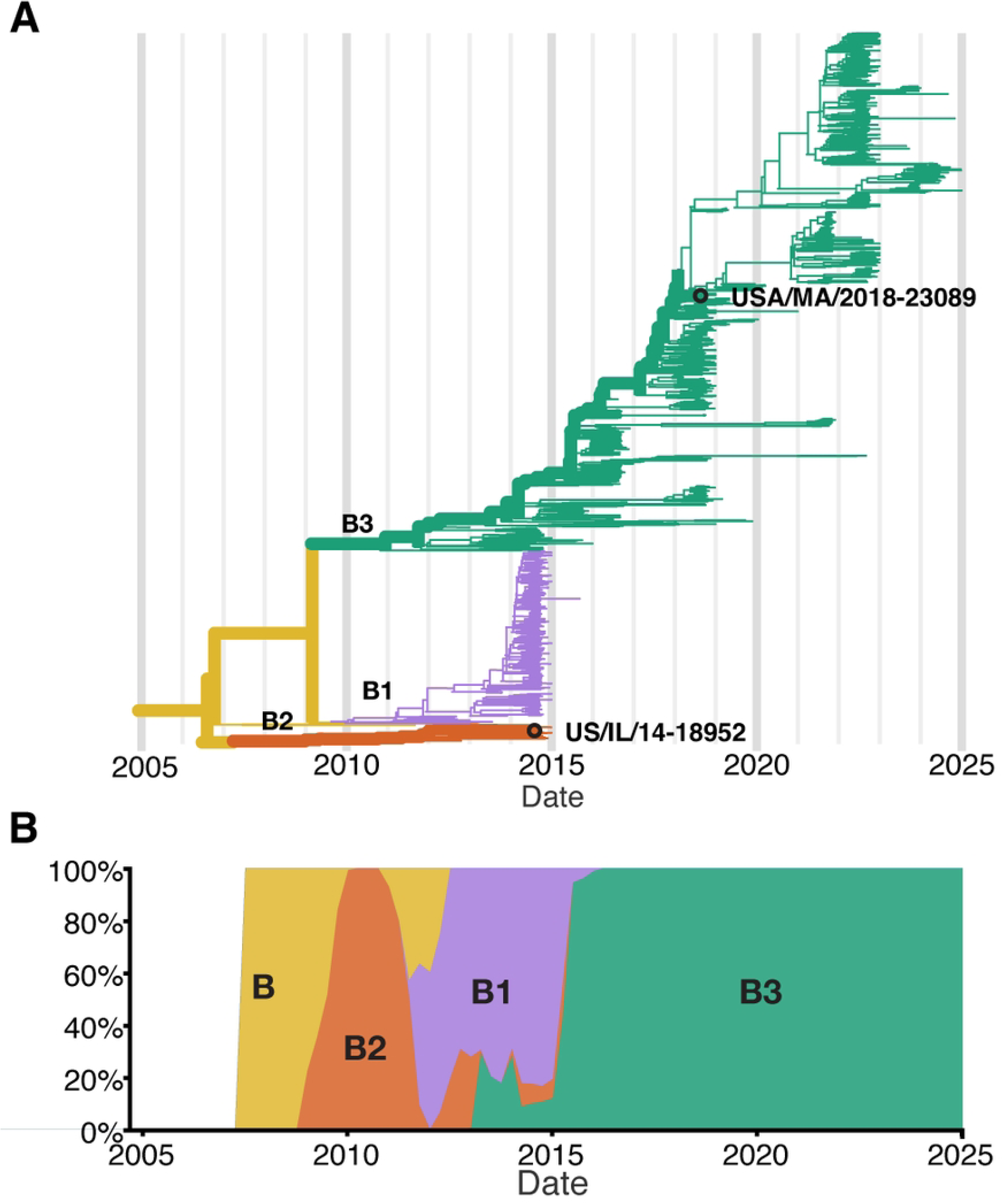
Nextstrain Phylodynamics of Enterovirus D68 Clade B. (A) Phylogenetic tree of publicly available EV-D68 Clade B sequences based on VP1 region (N = 1311 sequences; collection dates: April 2008 to December of 2024), highlighting the two strains used in this study - US/IL/14-18952 (18952) and US/MA/18-23089 (23089). (B) Temporal distribution of EV-D68 Clade B sequences, shown as the proportion of sequences per year.

## Results

### 24-days-old spinal cord organoids comprise diverse cell lineages

Previous characterization of marker gene and protein expression in hSCOs suggested the presence of several fully-differentiated cell types, including neuronal lineages, such as motor neurons and interneurons, and roof plate-like structures and neuroepithelium [21,25]. However, such analyses of gene expression may not identify minor cell types or developmental intermediates of specific cell lineages, and do not provide quantitative measures of relative cell abundance. Therefore, to better characterize the diversity and abundance of individual cell types in these organoids, we performed scRNAseq. We generated single-cell suspensions by dissociating pools of 12 hSCOs differentiated for 24 days, capturing between 8,000 and 10,000 cells per sample for sequencing (Fig 2A).

**Figure 2.**
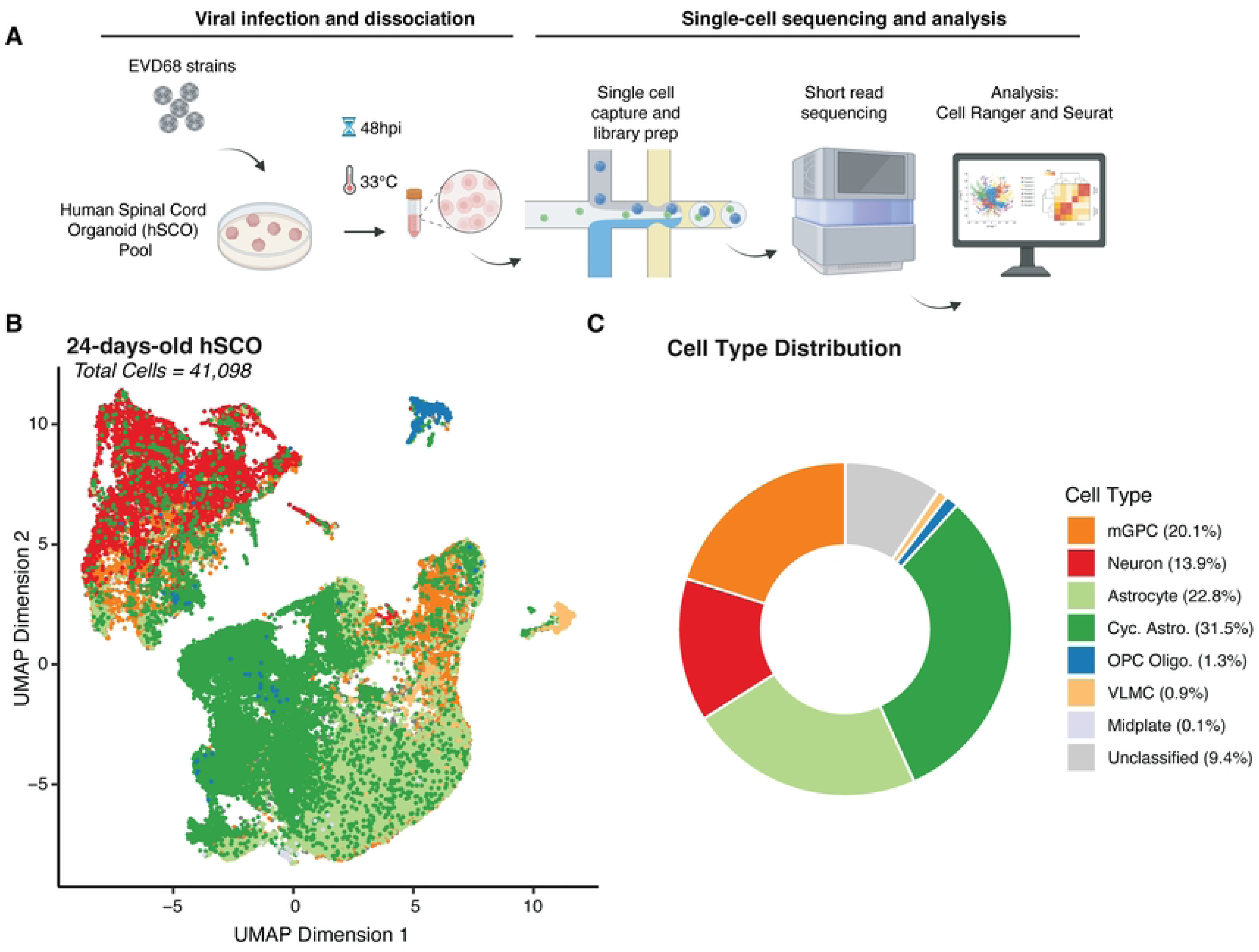
Cell type distribution in 24-day-old hSCOs. (A) Design of scRNAseq experiment. Pooled hSCOs (n=12) were infected with EV-D68 strains, US/IL/14-18952 (18952) or US/MA/18-23089 (23089) (10^5^ PFU/group) for 48 hours at 33°C, then dissociated mechanically and enzymatically. Cells were partitioned using the 10x NextGEM procedure and resulting libraries were sequenced using short-read sequencing. Flowchart created with BioRender. (B) Uniform Manifold Approximation Projection (UMAP) of individual cells identified from all the samples collected in this study colored by assigned cell type (41,098 cells). (C) Donut plot showing the relative abundance of cell types captured across all experiments.

Following scRNAseq, we used gene expression profiles from individual cells to perform cell type identification. We based our assignments on curated cell types from a previously published study of the developing human spinal cord [26] through “label transfer” [27] (Fig 2B-C). This analysis revealed that hSCOs exhibit a diverse cellular composition, encompassing neuronal lineages, along with astrocytes, oligodendrocytes and their progenitors (OPCs), glial populations such as midplate cells and multipotent glial progenitor cells (mGPCs), and vascular leptomeningeal cells (VLMCs), which contribute to blood-brain barrier integrity.

Astrocytes and cycling astrocytes were identified as the majority cell type in the 24-old day hSCOs (54.3%). We confirmed their cell identity by assessing expression of known astrocyte-specific markers, including *SOX9*, *FGFR3* [28,29] *and TOP2A* [30] (S1 Fig). Neurons comprised the next most common fully-differentiated cell type (13.9%), which we confirmed by examining expression of *MAP2* and *TUBB3* [31,32] (S1 Fig). In addition to differentiated cell types, mGPCs also made up a considerable proportion of the hSCO composition (20.1%) (Fig 2B-C, S2-A Fig). These proportions were consistent across replicate pools of hSCOs (S2-B Fig) of 24-days-old hSCOs. Fewer than 10% of cells were not classified as a specific cell type due to low predicted cell type score (Fig 2, “Unclassified”). These were not considered for subsequent analysis.

Focused reanalysis of the mGPC subset revealed five distinct clusters which may represent intermediate states in the differentiating organoid (S3-A Fig). Consistent with this interpretation, we have found the proportion of mGPCs are reduced in hSCOs in later development days (data not shown). mGPCs in clusters 0 and 4 display features of neural progenitors and early neuronal differentiation, with Cluster 0 exhibiting expression of genes linked to mature neuronal identity, while Cluster 4 shows expression patterns indicative of active neurogenesis. Among the expression differences, we identified *DCX* and *NEUROG1* in Cluster 0, and *NKX1-1* and *GATA2* in Cluster 4, reflecting their roles in early neural development [33–36]. Cells in Cluster 0 showed high expression of *NEUROD4* and *ELAVL3* [37–39], markers of mature neurons. In contrast, Cluster 4 displays a broader range of functions, including neurotransmitter synthesis (*GAD2* and *SLC32A1*) [40,41], and cell adhesion and signaling (*GPR83* and *CNTNAP5*) [42–44], indicating a more diverse cell population (S3-B Fig). Clusters 1, 2, and 3 are likely astrocyte progenitor cells, with diverse gene expression profiles related to stress response, lipid metabolism, cell signaling, extracellular matrix remodeling, DNA repair, cell cycle regulation, neural development, and cell polarity. Cluster 2 exhibited markers of proliferation, more consistent with cycling astrocytes.

### Two contemporary EV-D68 strains show distinct tropism in hSCOs

Upon infection with two distinct strains of EV-D68 (US/IL/14-18952 and US/MA/18-23089), we observed similar patterns of broad cellular susceptibility across the three main cell types identified within the hSCOs (Fig 3-A). Overall, we identified 144 viral RNA (vRNA)-positive cells out of 8,698 total cells in the 18952-infected organoids, representing 1.65% of the captured cell population. For strain 23089, we observed 205 vRNA-positive cells out of 15,805 total cells, corresponding to 1.3% of the captured cells. Although the proportion of infected cells is similar, the two strains exhibited distinct preferences for specific cell types. EV-D68-18952-infected cells were significantly enriched in neurons (based on permutation tests), whereas EV-D68-23089-infected cells were significantly enriched among cycling astrocytes and oligodendrocyte progenitor cells (OPC Oligo.) (Fig 3-B). Notably, these enrichments were also reflected in the number of viral RNA reads (vRNA) originating from these cell types. Although, of note, we observed significant heterogeneity in vRNA reads per individual cell (Fig 3-C). Together these observations suggest marked strain differences in host cell preference in the spinal cord, which may contribute to differential pathogenesis although the consequences are not clear from this observation alone.

**Figure 3.**
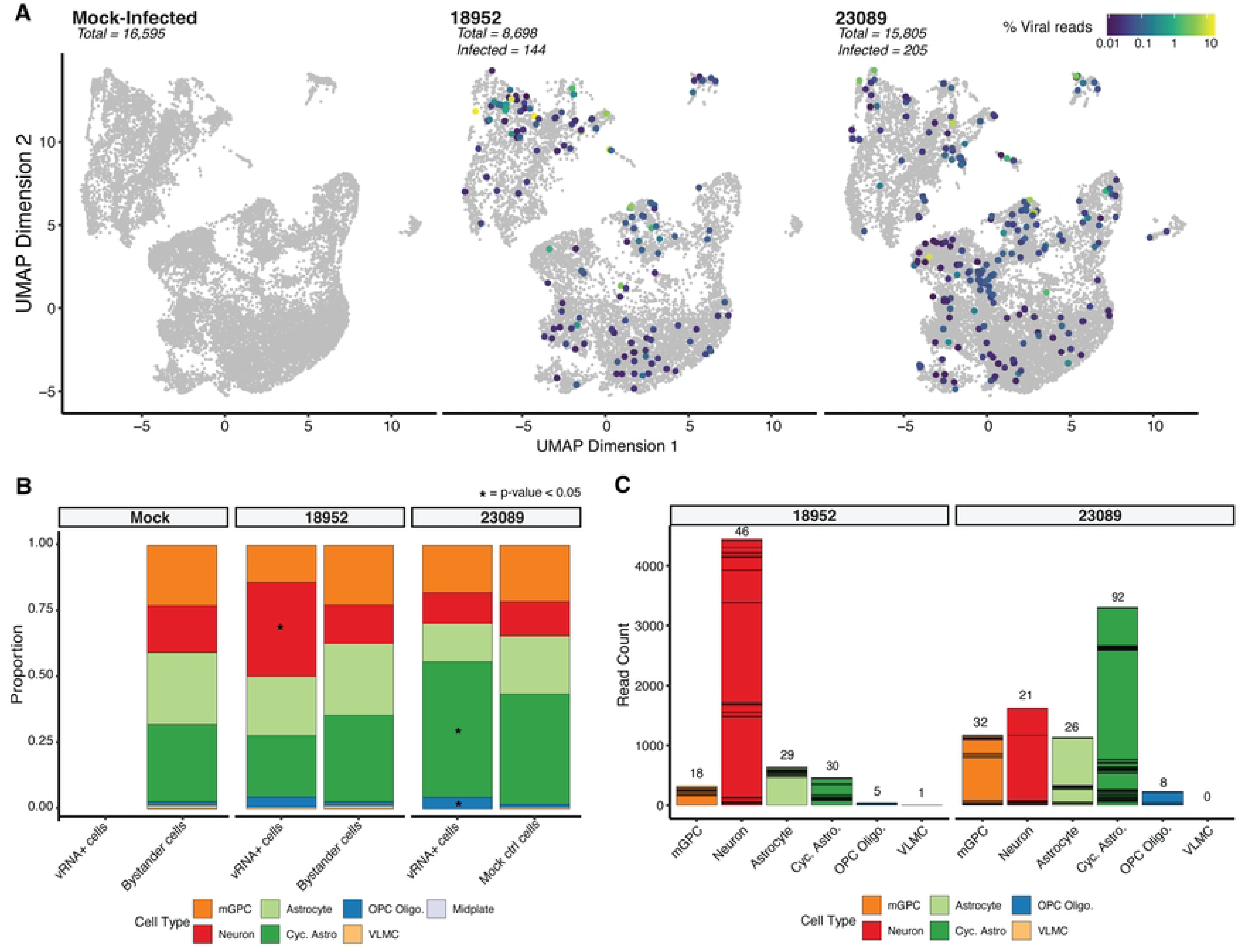
Infection profiling in hSCO infected with EV-D68. (A) UMAP of cellular transcriptional phenotypes in 24-day-old organoids infected with EV-D68 strains 18952 and 23089. Cells positive for viral RNA are shown colored by viral RNA content. (B) Bar plots displaying the proportion of vRNA+ cells vs. Bystander cells in various cell types per each condition, highlighting significant differences with an asterisk. (C) Stacked bar charts showing viral read counts in each cell type. The number of total stacked cells are indicated on the top of each bar.

### Strain-Specific Enrichment Analysis Reveals Distinct Responses to EV-D68 Infection

To address the distinct cellular and molecular responses triggered by two strains of EV-D68 (18952 and 23089), we performed a comprehensive enrichment pathway analysis on infected spinal cord organoids to identify strain-specific transcriptional programs. Due to the low proportion of vRNA-positive cells, we combined all cell types to compare gene expression differences based on the infecting strain. While this approach does not resolve cell type-specific changes, it allows for a more statistically robust comparison of strain-specific differences in host response. The normalized enrichment scores (NES) of significantly enriched pathways (adjusted p < 0.05) revealed three major functional clusters (Fig. 4A), capturing pathways where genes are primarily up-regulated (*Cluster 1*) or down-regulated (*Cluster 3*) in response to infection with either strain, or pathways primarily upregulated in response to 23089 infection relative to 18952-infected (*Cluster 2*). Notably, our analysis did not identify any pathways enriched only in the context of 18952 infection.

**Figure 4.**
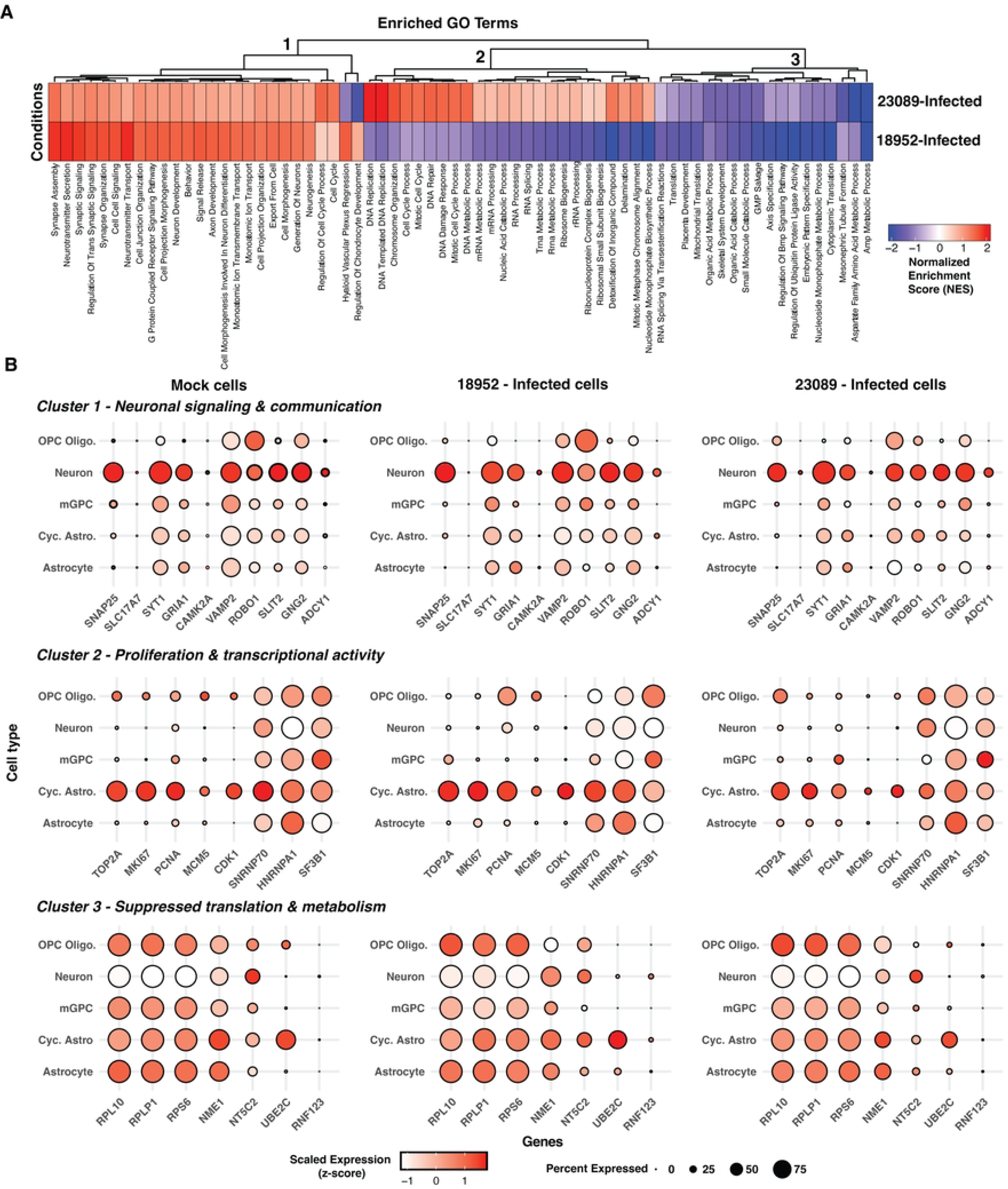
Strain-specific enrichment of host pathways in infected cells. (A) Heatmap of normalized enrichment scores (NES) from gene set enrichment analysis (GSEA) comparing two aggregated cell groups: 18952-Infected and 23089-Infected. Pathways shown represent all significantly enriched Gene Ontology Biological Process (GO-BP) terms (adjusted p < 0.05) including both positively and negatively enriched gene sets. Negative NES values (blue) reflect pathways more enriched in Mock cells relative to the infected group. (B) Dot plots showing the expression of key genes involved in neuronal signaling and communication, proliferation and transcriptional activity, and suppressed translation and metabolism processes across distinct cell types in hSCOs infected with EV-D68 strains 18952 and 23089. Dot size represents the percentage of cells expressing each gene, while color intensity reflects the average expression level, scaled (z-score) across all cells based on log-normalized values. Midplate and VLMCs were excluded from the Mock condition, as they were either absent from the infected groups or detected in only one of them (e.g., VLMCs: 1 cell in EV-D68-18952).

The most striking difference between the two strains is in Cluster 2, which corresponded to proliferative and biosynthetic transcriptional programs uniquely enriched in 23089-infected cells. This included pathways such as *DNA replication*, *chromosome organization*, *cell cycle*, and *ribosome biogenesis* (Fig. 4A). High NES values were observed for *DNA-templated DNA replication* (NES = 2.07), *DNA repair* (NES = 1.56), and *cell cycle process* (NES = 1.52). Despite 18952-infected cells showing slightly higher expression of cell cycle markers per cell (Fig. 4B), the enrichment in 23089 is likely driven by the larger number of infected cycling astrocytes, increasing the overall representation of these proliferative pathways. 23089-infected cells showed modest upregulation of RNA processing and splicing pathways across all infected cell types (e.g. *SNRNP70*, *HNRNPA1* and *SF3B1*) (Fig. 4B). These findings suggest that 23089 induces a transcriptionally-active and biosynthetically engaged state, possibly reflecting its preferential infection of cycling astrocytes.

Clusters 1 and 3 captured similarities in response to the two strains. Pathways identified in Cluster 1 included neuronal development and communication—such as *synapse assembly*, *axon development*, *neurotransmitter secretion*, *G protein-coupled receptor signaling*, and *neurogenesis*—which were positively enriched in both 18952- and 23089-infected cells, consistent with the neurotropic nature of EV-D68. However, strain 18952 showed higher enrichment for *synaptic signaling* (NES = 1.87 vs. 1.13) and *neurotransmitter transport* (NES = 1.95 vs. 1.13), suggesting a more robust neuronal response (Fig 4A). Cluster 3 encompassed pathways related to translation, metabolism, and developmental signaling, with 18952 showing stronger negative enrichment, including *cytoplasmic translation* (NES = –2.22), *AMP metabolic process* (NES = –2.26), and *ubiquitin ligase regulation* (NES = –2.30). While genes encoding ribosomal proteins (e.g. *RPL10*, *RPLP1* and *RPS6*) appeared downregulated in neurons from both infected and mock conditions (Fig 4B), broader suppression of metabolic homeostasis pathways was more pronounced in the infected cells. Together, these clusters suggest that both strains impact neuronal function and basal cellular processes, but with greater intensity in 18952-infected cells.

These transcriptional differences align with the distinct cellular tropism observed between the two EV-D68 strains. In organoids infected with 18952, neurons represented the major significantly enriched infected cell population, which is consistent with the prominent upregulation of synaptic signaling, axonal development, and neurotransmitter-related pathways. The robust enrichment of these neuronal processes suggests a direct viral impact on neurons, potentially contributing to neuronal dysfunction or degeneration. In contrast, 23089 predominantly infected cycling astrocytes and, to a lesser extent, oligodendrocyte progenitor cells (OPCs). This cell-type specificity is mirrored by the enrichment of DNA replication, cell cycle progression, and RNA processing pathways in 23089-infected cells—hallmarks of transcriptionally active, proliferative glial populations. Thus, the pathway signatures not only reflect divergent viral-host interactions but also underscore how strain-specific cellular targeting shapes the overall transcriptional landscape of infected organoids.

### Distinct Transcriptional Impacts of EV-D68 Strains Across Bystander Cell Types

Differential gene expression (DEG) analysis was performed on bystander cells stratified by cell type. A summary of the number of significant DEGs identified per cell type is presented in (Fig. 5A). This panel reflects DEGs filtered by both adjusted p-value < 0.05 and absolute log_2_ fold-change greater than 0.25, ensuring that the displayed genes represent biologically relevant transcriptional shifts. Significant DEG counts were observed for neurons, astrocytes, and cycling astrocytes, whereas OPCs did not show any DEGs that met these criteria. Accordingly, they were analyzed similarly to the vRNA-positive cells.

**Figure 5.**
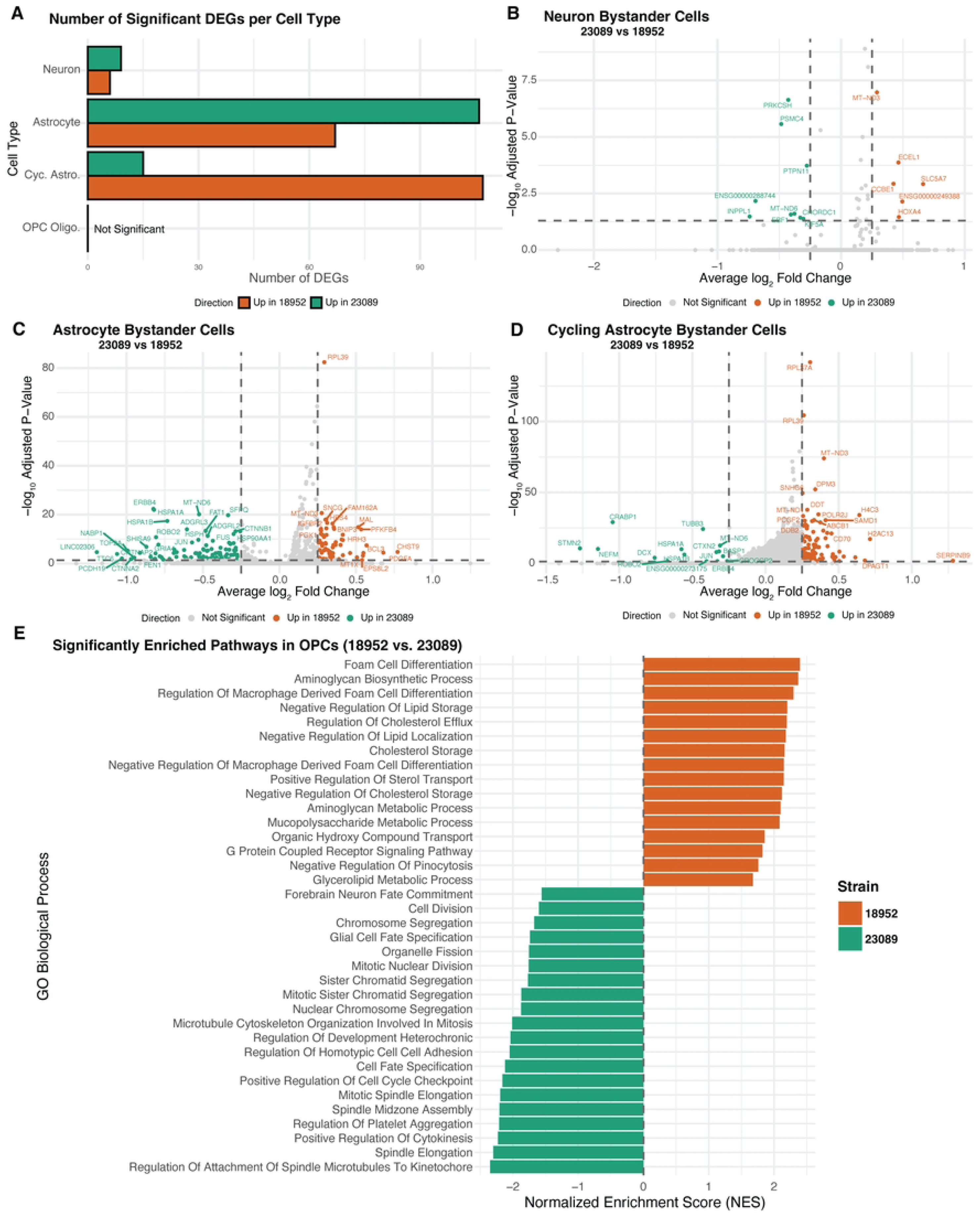
Strain-specific transcriptional responses in bystander cells. (A) Bar plot showing the number of significantly differentially expressed genes (DEGs) between EV-D68 strains 18952 and 23089 across bystander neurons, astrocytes, cycling astrocytes, and OPCs. DEGs were defined as adjusted p-value < 0.05 and log₂FC > 0.25. B–D). Volcano plots displaying DEGs between 23089- and 18952-exposed bystander cells in neurons (B), astrocytes (C), and cycling astrocytes (D). Genes upregulated in strain 23089 are shown in green, and those upregulated in strain 18952 are in orange; non-significant genes are shown in gray. Dashed lines indicate significance thresholds (adjusted p = 0.05 and log₂FC = ±0.25). (E) Bar plot showing normalized enrichment scores (NES) from GSEA of OPC Oligo. bystander cells, comparing transcriptional profiles between strains 23089 and 18952. Despite the absence of individual DEGs meeting significance, GSEA revealed pathway-level differences. Bars are colored by the direction of enrichment: green for pathways enriched in 23089 and orange for pathways enriched in 18952.

The transcriptional response in bystanders varied notably according to both cell type and viral strain. Among the different cell types, neurons demonstrated a short list of DEGs, while astrocytes and especially cycling astrocytes showed a more pronounced transcriptional response. Neurons and astrocytes displayed a greater number of DEGs in response to the 23089 strain, suggesting a more pronounced impact of this viral strain on these cell populations. In contrast, cycling astrocytes exhibited more DEGs associated with the 18952 strain. However, the magnitude of transcriptional changes differed between strains: DEGs associated with 18952 tended to have more modest fold changes, whereas 23089 DEGs spanned a broader range of fold changes (Fig 5D). These patterns indicate that both the extent and intensity of the bystander transcriptional response differ depending on cell type and viral strain.

Neurons displayed a comparatively limited bystander transcriptional response, with fewer differentially expressed genes than astrocytes or cycling astrocytes. In the 18952 condition, upregulated genes included *SLC5A7*, essential for acetylcholine synthesis and synaptic transmission [45], which is consistent with the cholinergic nature of spinal motor neurons. *ECEL1*, implicated in motor neuron axon development [46], was also elevated. Other transcripts, such as *HOXA4* and *CCBE1*, related to positional identity and extracellular matrix organization [47,48], suggest a role in preserving neuronal identity and structure.

In contrast, 23089-exposed neurons upregulated stress-responsive and protein quality control genes. These included *PTPN11* [49], *CHORDC1*, and *PRKCSH*, associated with *MAPK* signaling, protein folding, and ER stress [50,51]. Increases in *MT-ND6* and *PSMC4* expression, involved in mitochondrial respiration and proteasomal degradation [52,53], may reflect compensatory responses to maintain homeostasis (Fig. 5B). These trends suggest that while 18952 maintains neuronal identity, 23089 induces a mild stress-adaptive transcriptional state.

Astrocytes exhibited a broader transcriptional response than neurons, with distinct gene expression patterns between strains. In the 18952 condition, upregulated genes included *CHST9* and *PDGFA*, involved in extracellular matrix remodeling and glial signaling, alongside *HRH3* and *SLC2A3*, linked to histaminergic modulation and glucose transport. Additional genes such as *BCL3, PFKFB4*, and *MT1X* suggested mild activation of redox regulation and metabolic homeostasis. Together, these changes reflect a modestly reactive but structurally supportive astrocyte state. In contrast, astrocytes exposed to the 23089-strain upregulated a distinct set of genes, including *HSPA1A*, *CNTNAP2*, *PCDH19*, and *ROBO2*, which are involved in synaptic regulation, axon-glia interaction, and cellular stress responses. The transcriptional profile also included *ERBB4*, *LINGO2*, and *TOP2A*, further implicating altered signaling and adhesion programs. These differences suggest that 23089-exposed astrocytes enter a more transcriptionally active state, potentially affecting communication with the neuronal environment. The overall transcriptional divergence reinforces the strain-specific modulation of astrocyte identity in the bystander context (Fig. 5C).

Cycling astrocytes displayed a unique transcriptional profile distinct from both neurons and non-cycling astrocytes, marked by broader identity and functional shifts between strains. In the 18952 condition, upregulated genes included *SERPINB9*, *DPAGT1*, and *LOX*, along with histone-associated transcripts such as *H2AC13*, *H4C3*, and *HIST1H1B*. These changes support a transcriptional program favoring immune regulation, extracellular matrix organization, and cell cycling. 23089-exposed cycling astrocytes showed a marked shift in identity, with upregulation of neuronal and stress-related genes including *STMN2*, *NEFM*, *DCX*, and *CRABP1*. Additional activation of *HSPA1A*, *JUN*, and *ERBB4* suggested elevated stress signaling and cytoskeletal remodeling, consistent with transcriptional dysregulation. The emergence of neuronal lineage markers in this glial progenitor population may reflect stress-induced unexpected activation of developmental genes not typically active in astrocyte progenitors [54].

Notably, this mixed transcriptional phenotype in cells predicted to be cycling astrocytes may be linked to the fact that these cells are also the primary targets of infection by the 23089 strain. Even in bystander cells, signaling cues from nearby infected cells could perturb their transcriptional program and disrupt normal progenitor dynamics. Compared to non-cycling astrocytes, which maintained more canonical glial features, the 23089-exposed cycling astrocytes exhibited broader deviations in transcription. These distinctions highlight the unique vulnerability of proliferative astrocyte populations to strain-specific viral influence (Fig. 5D).

In OPC Oligodendrocytes, pathway enrichment analysis revealed distinct transcriptional programs between strains, even in the bystander population. Due to the relatively low number of OPC Oligo. in this group, we adopted the same strategy used for infected cell analysis, performing enrichment pathways analysis to enable broader biological interpretation since no gene by itself stands out from our DEG analysis. OPCs exposed to the 18952-strain showed upregulation of pathways associated with lipid metabolism and extracellular matrix organization, including *foam cell differentiation*, *regulation of cholesterol efflux*, and *aminoglycan biosynthetic process*. These processes are important for membrane dynamics and maintenance of oligodendrocyte identity, suggesting that in the 18952 condition they may retain a more metabolically stable and structurally supportive state.

In contrast, OPCs exposed to the 23089-strain were enriched for pathways related to cell cycle regulation (*mitotic spindle elongation, cytokinesis*), microtubule organization, and developmental signaling, including *glial cell fate specification*. These transcriptional signatures are consistent with a disturbed cellular state, which could potentially reflect direct infection-related stress or dysregulation, as we observe for cycling astrocytes. This is in line with the fact that OPCs, alongside cycling astrocytes, represent a significant infected population in the 23089 condition. The enrichment of mitotic and developmental signaling programs may indicate that viral presence disrupts the normal proliferative or lineage-committed state of these glial progenitors. Together, these findings suggest that OPCs respond to 23089 exposure with transcriptional shifts that may compromise their homeostatic roles (Fig. 5E).

Finally, we noted a lack of altered expression of innate immune genes, particularly interferon stimulated genes (ISGs), in our organoid model. Further analysis revealed we could detect expression of genes known to be regulated by Type I, II, and III interferon in both infected and uninfected organoids suggesting the lack of induction was not due to limited detection in our sequencing (Supp. Fig. 4). Although, more robust immune responses may occur at later time points in infection (beyond 2dpi), this lack of ISG induction is consistent with previous studies highlighting the effectiveness of viral innate immune suppression in EV-D68 infection, through protease cleavage of innate immune signalling factors and the action of other viral proteins [55,56].

## Discussion

Our analysis of 24-day old organoids showed a diverse cell composition, with both neuronal and glial lineages, consistent with previous IF marker analysis [25]. While we identified neurons, mGPCs, OPCs, VLMCs, and more, the most common cell type in the hSCO at this age was cycling astrocytes and astrocytes (54.3%). Given the relatively nascent nature of the hSCO model, this majority may be due to astrocytes’ vital role in developing and maintaining neuronal functions throughout spinal cord development [57–60]. mGPCs (20.1%) also have the capacity to differentiate into both astrocytes and oligodendrocytes so cellular composition in more mature hSCO may shift [61]. Although this cell population has not yet committed to a differential pathway, they are still more specialized and committed than a broad progenitor cell. 24-day old organoids allow for the growth and development of CNS cell types without compromising on overall cell viability [21]. The abundance of CNS cell types present and interacting in hSCO allow for us to better understand EV-D68 tropism and dynamics in the complex human spinal cord.

To understand which CNS cell-types are infected by EV-D68, we performed scRNA-seq on hSCO infected with contemporary EV-D68 strains US/IL/14-18952 and US/MA/18-23089. We found that while both strains had the capacity to infect neuronal and glial cell lineages, 18952 preferentially infected neurons while 23809 preferentially infected cycling astrocytes and to a lesser extent, oligodendrocyte precursor cells. These differences extended to bystander cells — i.e., uninfected cells within infected hSCOs — where we observed more DEGs in 18952-bystander cycling astrocytes, and even larger fold changes in 23089-bystander neurons and astrocytes. These findings indicate that glial cell populations such as astrocytes and oligodendrocytes play an important role in EV-D68 associated AFM pathogenesis, varying amongst different strains and clade classification. Previous studies have implicated spinal cord neurons in EV-D68-induced paralysis, but our results suggest additional cell types are infected and transcriptionally altered, potentially playing unidentified roles in EV-D68 pathogenesis [13–17].

Astrocytes participate in both innate and adaptive immune system, such as regulating the release of cytokines and chemokines, thereby leading to antigen presentation that lead to recruitment of helper T-cells in pathogen or damage-affected areas of the CNS [62–66]. Several of the differentially expressed genes identified in bystander astrocytes and cycling astrocytes, particularly in the 23089 condition, are associated with stress signaling pathways. These changes, combined with the disruption of glial identity in 23089-exposed cycling astrocytes, may create an environment more permissive to immune cell recruitment. In contrast, the transcriptional profile in 18952-exposed astrocytes suggested a more limited activation of broad immune-modulatory programs, although cycling astrocytes did express immunoregulatory markers such as *CD70* and *SERPINB9*, potentially reflecting a more localized or cell-type stress response rather than a widespread inflammatory activation. These findings raise the possibility that differential modulation of innate immune signaling by each strain could further shape their pathogenic outcomes *in vivo*. Since we do not know yet to what degree the immune response mediates EV-D68-associated AFM, further studies are needed to assess astrocyte function and subsequent immune response during EV-D68 infection. Additionally, because oligodendrocytes form myelin to aid neuron conductivity and communication, its functionality upon EV-D68 infection should also be assessed [67,68].

We also identified strain-specific differences in cellular tropism within hSCO, indicating that there has been a change in EV-D68 viral infection dynamics from 2014 to 2018. EV-D68-18952 strain primarily infected neurons, while 23089 preferentially infected cycling astrocytes and, to a lesser extent, OPCs. Together, these results suggest that the two strains may induce distinct forms of cellular vulnerability — 18952 through direct neuronal targeting and 23089 through glial destabilization. While further studies will be needed to determine the long-term impact of these responses, the transcriptional divergence observed across cell types points to fundamentally different modes of pathogenesis. These differences may be broadly attributed to clade specific pathogenic differences between 18952 (B2) and 23089 (B3) or be pathogenic variations between two specific isolates. While both viruses are expected to cause the same clinical paralysis phenotype, the differences in tropism may have caused varying mechanisms of pathogenesis and we are not able to assess clinical differences between these two isolates.

Additional strain specific differences are reflected in the transcriptional impacts in bystander cells. Astrocytes and cycling astrocyte bystander cells had significantly more transcriptional changes compared to neuronal bystanders, with 23089 having more impact on astrocyte and neuron bystanders and 18952 having more of an impact on cycling astrocyte bystanders. Many of the upregulated genes in 23089 bystander cells were cellular stress indicators, primarily found in astrocytes. These cellular responses may also indicate a downstream loss in communication or homeostasis in neurons, as astrocytes are integral to maintaining neuronal function and stability. 23089 had a larger fold transcriptional shift in bystander cells compared to 18952. Although these are only two isolates, these differences may point to clade- or outbreak-specific variation in EV-D68 pathogenesis. The higher number of AFM cases in 2018 compared to 2014 could reflect such viral differences, but may also stem from multifactorial causes, including co-circulation of other enteroviruses (e.g., EV-A71) or improved case detection and reporting [69].

The OPC bystander population did not have any significant shifts, 18952 exposed OPCs were more stable and 23089 exposed OPCs had evidence of a stress response that interfered with glial cell differentiation. EV-D68 may be interfering with oligodendrocyte differentiation and maturation in the hSCO and thus prevent a fully functioning oligodendrocyte population. Considering the hSCO population of oligodendrocytes were precursor cells and still differentiating at the time of EV-D68 infection, cellular response may differ in mature oligodendrocytes. To better understand oligodendrocyte’s response to EV-D68 infection, studies in more aged hSCO will be necessary.

While hSCO scRNAseq has allowed us to further understand EV-D68 infection dynamics in the spinal cord, it does have its limitations. Since hSCO cells are relatively immature and represent a developing human spinal cord, EV-D68 infection dynamics may differ in hSCO to that of a child. Despite this limitation, its multicellular complexity and physiological relevance for AFM has and can let us learn more about EV-D68 pathogenesis in the CNS. This limitation in hSCO also lends itself to be an advantage when looking at enteroviruses that target neonates in order to better understand their tropism and pathogenesis.

Another limitation in our study is the relatively low number of infected cells identified for both EV-D68 strains. Despite leveraging an aggregated, bulk analysis approach to enhance statistical power, the small fraction of virus-positive cells could potentially limit the detection of more subtle transcriptional changes and low-abundance cell populations responding to infection. This may be particularly relevant for less abundant cell types, such as OPCs. Moreover, oligodendrocytes are known to be particularly fragile during tissue dissociation, potentially leading to underrepresentation in the final dataset. Additionally, the low number of infected cells observed could also reflect loss during dissociation or capture, particularly if infected cells were damaged and lysed during infection or processing.

Despite these limitations, our study provides valuable insights into the cellular and molecular landscape of EV-D68 infection in human spinal cord organoids, revealing distinct cell-type tropism for two contemporary strains. By integrating pathway enrichment analysis with single-cell transcriptomic profiling, we demonstrate that EV-D68-18952 exhibits a pronounced preference for neurons, driving synaptic signaling and axonal development pathways, while EV-D68-23089 predominantly targets cycling astrocytes, triggering transcriptional programs associated with cell cycle progression and RNA processing. These findings represent a step forward in understanding the strain-specific interactions of EV-D68 with neural populations, which may have implications for viral spread and neuropathogenesis. Furthermore, our use of spinal cord organoids as a model provides a physiologically relevant system to dissect host–virus interactions at single-cell resolution, underscoring the utility of this platform for studying neurotropic viruses.

## Materials and Methods

### Viruses and cells

EV-D68 US/IL/14-18952 (CDC) and US/MA/18-23089 (CDC) strains were propagated using HeLa cells incubated at 33°C and 5% CO_2_ and purified using sucrose-cushion as previously described [70]. These stocks were previously sequenced to confirm their identity with VP1 primers^3^.

HeLa 7b (ATCC, CCl-2) cells were maintained in MEM medium (ThermoFisher, 11095-072), supplemented to contain 5% FBS (Phenonix Scientific, PS-100), 1% penicillin/streptomycin (Corning, 30-002-Cl), and 1% NeAA (Corning, 25-025-Cl). Cells were grown at 37℃ and 5% CO2.

Human iPSC line SCTi003A (STEMCELL Technologies, 200-0511) was maintained in mTeSR^TM^ Plus medium (STEMCELL Technologies, 100-0276), supplemented with 10 µM Y-27632 (Tocris, 1254). They were seeded and passaged in flasks coated with 150 µg/mL Cultrex (R&D Systems, 3434-005-02). The 3-DiSC hSCO were propagated and differentiated as described in Aguglia et al. [21] for up to 24 days.

### hSCO infections and dissociations

hSCOs were infected in pools of 12 organoids each and inoculated with virus at 10^5^ PFU/pool. After 1 hour of incubation at room temperature, hSCOs were washed 3X with PBS and moved to new wells before incubation with fresh medium. No further media changes were performed for the rest of the experiment. The EV-D68 infected pool was incubated at 33°C for 48 hours post infection (hpi). After 48hpi, the pools were dissociated to a single-cell suspension with Accumax (Invitrogen). They were incubated for 15 minutes in the water bath at 37℃, gently mixed, and then proceeded with the proposed 10X Genomics protocol (Cell Preparation Guide - CG00053 Rev C). Once the cell’s concentration was achieved the cells were moved to ice.

### Single cell RNAseq cDNA library generation

All samples were calculated to achieve ∼5000-8000 targeting cells in the single-cell preparations using Chromium Next GEM Single Cell 5’ standard kit. For the preparation of the cDNA and sequencing library generation, we followed the instructions from the user guide Chromium Next GEM Single Cell 5’ Reagent Kit v2 (Dual index). All other steps were followed to produce cDNA and subsequent Illumina sequencing library for single cell sequencing. The illumina library preparation was submitted to quality control in the TapeStation D1000 high sensitivity for size distribution and DNA concentration was measured by Qubit High Sensitivity dsDNA kit. The molar concentration of the libraries were determined and the samples were diluted for sequencing according to illumina sequencing protocol. We aimed to sequence each library to achieve ∼50,000 reads per cell.

### CellRanger

CellRanger v7.0.0 was used [71] to align reads against a composite genome, which encompassed both the GRCh38 human reference genome and the *Enterovirus* genome corresponding to the specific strain identity of the sample. Feature-barcode matrices were generated using GENCODE v44 GRCh38 gene models and the viral strain’s genome as a single ORF. Default parameters of the ‘cellranger count’ function were used.

### Seurat

Seurat v5.0.2 was used [72]. CellRanger gene counts were made compatible across different strains by setting the name of the gene encompassing all viral reads for each sample to ‘Viral-Gene’. Cells with mitochondrial gene expression > 25% or detected gene counts < 1000 were removed. Gene counts of filtered cells were normalized using SCTransform, while specifying the ‘vars.to.regress’ parameter to the ‘S.Scores’ and ‘G2M.Scores’ obtained from the ‘CellCycleScoring’ function, utilizing Seurat’s ‘cc.genes’ cell cycle gene list. Reciprocal PCA integration analyses were then performed to generate integrated datasets for EV-D68. Dimensionality reduction of the integrated dataset was performed using the ‘RunPCA’ function and ‘RunUMAP’ function using the top 30 principal components. Cell cluster analysis was performed using the ‘FindNeighbors’ function using the top 30 principal components and the ‘FindClusters’ function using a ‘resolution’ of 0.1. Cell type annotations were predicted using cell label transfer with Seurat’s FindTransferAnchors, TransferData, and AddMetaData functions using a previously annotated spinal cord scRNAseq dataset [26] as our reference, considering “true” cell types above 0.5 prediction.score.max. The cells that were below this threshold were assigned as NA or Unclassified.

### Assignment of infection status

Viral read percentages were calculated as the fraction of total UMIs per cell mapping to the viral gene (i.e. ‘Viral-Gene’) and were computed using the ‘PercentageFeatureSet’ Seurat function. To classify cells as ‘Infected’ or ‘NonInfected’, a Poisson test was applied to these percentages using the ‘estimateNonExpressingCells’ function from the SoupX R package [73], using an FDR of 0.05. This function accounts for ambient RNA contamination in the sample, which was estimated using SoupX’s ’autoEstCont’ function with ‘tfidfMin’ set to 1.0 and ‘soupQuantile’ set to 0.9.

### Permutation-based enrichment analysis

To assess whether specific cell types were significantly enriched in infected (vRNA+ cells) versus non-infected (Bystander cells) conditions, we performed a permutation test on cell count distributions across infection states. For each viral strain, we computed contingency tables comparing observed cell type frequencies across infection status (FDR < 0.05). To establish a null distribution, we randomly permuted cell type labels 10,000 times while preserving the infection status labels and recomputed the contingency tables for each iteration. Median values and 95% confidence intervals (2.5th and 97.5th percentiles) were derived from the permuted distributions. Observed counts exceeding the upper confidence interval were considered significantly enriched (p value < 0.05). Enrichment scores were calculated as the ratio of observed to permuted median counts for each cell type and condition. Analyses were conducted in R using the *data.table*, *Seurat*, and *SeuratObject* packages.

### Differential gene expression analysis

Differential gene expression (DEG) analysis was conducted using the *FindMarkers* function in Seurat v5. Two complementary approaches were applied. First, a pseudo-bulk-style analysis was performed by grouping cells according to their combined infection status and viral strain (e.g., “18952_Infected” vs. “18952_NotInfected”), enabling comparisons across aggregated conditions. This approach was used because the number of infected cells was insufficient for robust stratification by cell type. Second, a cell-type-specific DEG analysis was performed using only bystander cells, focusing on neurons, astrocytes, cycling astrocytes, and oligodendrocyte lineage cells (OPC Oligo.) to compare transcriptional responses between EV-D68 strains 18952 and 23089. All analyses were conducted using the RNA assay, with normalization via *NormalizeData*. DEG identification used a minimum expression threshold of 10% (min.pct = 0.1), no log fold-change cutoff (logfc.threshold = 0), and a minimum of 50 cells per group. Fold changes were calculated as log2-transformed values, and significance was assessed using Bonferroni-adjusted p-values. Genes with adjusted p value < 0.05 and | log₂FC | > 0.25 were considered significantly differentially expressed. Visualizations, including volcano plots and DEG count barplots by cell type and direction, were generated using *ggplot2*, *ggrepel*, and *patchwork*.

### Pathway enrichment analysis

Gene set enrichment analysis (GSEA) was performed using the *fgsea* package to identify biological pathways enriched in infected conditions or specific cell types, even in the absence of significantly differentially expressed genes. For certain DEG comparisons, particularly those involving low-abundance populations (e.g., infected cells or bystander OPC Oligo.), no genes met the adjusted p-value threshold for significance. In these cases, the full ranked DEG lists based on average log2 fold change were used as input for GSEA, enabling the detection of coordinated pathway-level shifts. Ranked gene lists were generated from pairwise comparisons of infection conditions (e.g., 18952-Infected vs. Mock) and bystander cell types (e.g., 23089 vs. 18952 within OPC Oligo.), and enrichment was computed using 100 million permutations for robust estimation. Gene sets were sourced from the Gene Ontology Biological Process (GO-BP) category via *msigdbr*. Normalized enrichment scores (NES) were computed for each condition, and pathways with adjusted p-values < 0.05 were considered significantly enriched. Heatmaps of NES values across conditions were generated using *ComplexHeatmap*, and selected pathway comparisons were visualized with *ggplot2* and *ggrepel*.

## Data Availability

Data from the scRNAseq analysis is deposited in Gene Expression Omnibus (GEO) (Accession number GSE292051).

## Supporting Information

**Supplementary Figure 1.** UMAP ‘FeaturePlot’ showing Log-normalized expression of key marker genes of astrocytes (*SOX9* and *FGFR3*), cycling astrocytes (*TOP2A*) and neurons (*MAP2* and *TUBB3*).

**Supplementary Figure 2.** Cell type characterization in hSCOs. (A) UMAP facets of all cell types in 24-day old hSCO. (B) Donut plots showing the relative frequency of cell types in hSCOs infected with each EV-D68 strain and mock-infected controls.

**Supplementary Figure 3.** (A) UMAP showing the cellular transcriptional phenotypes of mGPC clusters in 24-days old hSCO. (B) Heatmap of top 20 marker genes that distinguish each cluster. The heatmap displays scaled expression values (z-scores) of the top 20 marker genes per cluster, calculated using Seurat’s ScaleData function. Values are centered and scaled per gene across all cells, such that 0 represents the mean expression and ±2 corresponds to approximately two standard deviations above or below the mean. This highlights relative over- or underexpression patterns across clusters.

**Supplementary Figure 4.** Scatter Plots comparing of the proportion of cells expressing ISGs induced by Type I, II, and III interferons, and Volcano plots comparing the fold change in expression and the significance, as -log_10_(adjusted p-value), for comparisons of (A) Mock vs. 18952 Bystander cells, (B) Mock vs. 20892 Bystander cells, (C) 18952 Bystander cells vs. 20892 Bystander cells.

